# Constructing high-density linkage maps with MapDisto 2.0

**DOI:** 10.1101/089177

**Authors:** Christopher Heffelfinger, Christopher A. Fragoso, Mathias Lorieux

**Affiliations:** Department of Molecular, Cellular, and Developmental Biology, Yale University, New Haven, CT, USA; DIADE Research Unit, Institut de Recherche pour le Développement (IRD), Montpellier, France; Rice Genetics and Genomics Laboratory, International Center for Tropical Agriculture (CIAT), Cali, Colombia

**Keywords:** Genotyping by sequencing, High-density linkage map, Segregation distortion, Single-nucleotide polymorphism, Quantitative trait locus

## Abstract

We present the second major release of MapDisto, a multi-platform, user-friendly computer program for constructing genetic linkage maps of experimental populations. This version includes several new major features: (i) handling of very large genotyping datasets like the ones generated by genotyping-by-sequencing (GBS); (ii) direct importation and conversion of Variant Call Format (VCF) files; (iii) detection of linkage, *i.e.* construction of linkage groups in case of segregation distortion; (iv) data imputation on VCF files using a new approach, called LB-Impute. These features operate through inclusion of new Java modules that are installed and used transparently by MapDisto; (v) QTL detection *via* a new R/qtl graphical interface.

The program is available free of charge at mapdisto.free.fr.

## Introduction

MapDisto is a user-friendly computer program for construction of genetic linkage maps (Lorieux 2012). It has been used in numerous publications to construct maps of various plant, fungi or animal organisms^1^. In this note, we present a new implementation that can handle very large marker datasets obtained by genotyping-by-sequencing (GBS) (Elshire *et al.* 2011), and build linkage groups more efficiently in case of segregation distortion.

With the advent of genotyping techniques based on re-sequencing of segregating progeny genomes, it is now possible to generate extremely dense genetic linkage maps (He *et al.* 2014). GBS is one of the most promising among this family of methods (Elshire *et al.* 2011; Heffelfinger *et al.* 2014). The increasing size of datasets due the growth of GBS approaches in genetics research requires improvements in computation efficiency, both in terms of speed and use of random-access memory (RAM). GBS datasets typically contain thousands or tens of thousands of markers, generally single-nucleotide polymorphisms (SNP). This makes certain steps of linkage map construction quite challenging when it comes to mapping programs running on desktop computers. For example, determining linkage groups requires the calculation of *m*(*m* – 1)/2 statistical tests, e.g. recombination fractions or LOD scores. With a dataset comprised of 50,000 markers, this turns to 1,249,975,000 independent calculations, which corresponding matrix needs ~10 GB (gigabyte) to be stored in RAM if encoded as double-precision variables.

Another issue in linkage mapping is segregation distortion (SD). SD is defined as the deviation of parental allelic frequencies from their expected values in a segregating population. For example, in a recombinant inbred line (RIL) population, each parental allele has an expected frequency of 0.5. Extreme segregation distortion can affect the analysis of linkage, that is, the construction of linkage groups (LG) from a set of markers randomly sorted (Lorieux *et al.* 1995a; Lorieux *et al.* 1995b). Obviously, this step of LG construction isn’t necessary for species that benefit from a reference genome sequence (RefSeq), although even in datasets for these species it is important to verify correspondence between the physical and the genetic maps. For those species that do not have a high-quality RefSeq, it is crucial to be able to build reliable linkage groups, for example, if high-density genetic mapping is used to help organizing the contigs and scaffolds to build a RefSeq (Ariyadasa *et al.* 2014). Formulas to determine presence of linkage in presence of SD have been developed (Lorieux et al. 1995a; Lorieux et al. 1995b; Zhu et al. 2007). However, we found that these formulas do not take all possible situations into account. New algorithms and formulas are thus necessary to build high-confidence LGs.

As next generation sequencing becomes a dominant method for genotyping populations, especially through reduced representation technologies such as Genotyping-by-Sequencing and Restriction-Site Associated DNA Markers, new computational methodologies are required to ensure high quality data. Two issues are endemic to these datasets: missing and erroneous data. Missing data results when sequencing coverage is insufficient to recover marker calls at all sites. Erroneous data may result from a variety of sources ranging from sequencing to data processing. Another type of error, false homozygosity, occurs from incomplete allele recovery at a heterozygous site. This type of error is common in low-coverage sequences datasets for highly heterozygous populations, but may also affect inbred populations, such as RILs at a lower rate. Novel imputation algorithms that can not only impute missing data, but also resolve erroneous data are therefore required.

## Main new features

MapDisto now offloads a variety of operations to an included “MapDistoAddons.jar” Java package. The offloaded operations include Variant Call Format (VCF) file conversion, linkage group calculation, and data imputation. Similarly, data imputation and correction are performed by LB-impute (Fragoso et al. 2016) through the “LB-Impute.jar” Java module. QTL detection is performed via a new graphical interface, which runs R/qtl commands in the background.

### Large dataset handling and speed

The maximum dataset size depends on the amount of available memory. On modern desktop computers equipped with 16 or 32 GB of RAM, hundreds of thousands of markers could theoretically be processed. In the previous version (v.1.7) based on Visual Basic for Applications (VBA) 32-bit code, the maximum dataset that MapDisto could handle was about ~8,900 or ~16,000 markers on the Microsoft Windows and Apple OS X platforms, respectively, and ~32,000 on the 64-bit version of Microsoft Windows. This was due to internal limitations for array size of the VBA virtual machine. We also observed a 20–60x increase in computation speed – e.g., LOD scores or recombination fractions – compared to v.1.7.

As an example, we were able to process a rice population consisting of 181 RIL and scored with 44,398 GBS markers on a microcomputer equipped with as few as 4 GB of RAM and running OSX 10.10.5 and Excel 2011. It took only ~10 minutes to compute a complete LOD score and recombination fraction matrix and to determine the linkage groups. This is achieved thanks to implementation of multi-threading in the Java module.

### VCF conversion

MapDisto can now convert v.4.1 VCF files into MapDisto format and automatically import the converted file. If parents’ identifiers in the VCF file are specified, A (homozygous parent 1), B (homozygous parent 2) and H (heterozygous) alleles will be assigned to calls based on them. If parents’ identifiers are not specified, alleles at a given marker will be assigned based on the first allele observed in the dataset by the algorithm.

### Marker and individual filtering

Recombinant populations contain a limited number of recombination breakpoints. In highly-saturated genotyping experiments, like GBS assays, markers often completely co-segregate and the information they provide about recombination is thus redundant. The ‘Filter loci’ identifies the minimum number of genetic markers that provides the full recombination information in the population.

A new ‘Compute statistics on genotypes’ allows filtering of individuals of number of recombination breakpoints (or transitions), percentage of missing data, and percentage of genotypic configuration (homozygous parent 1 or 2, and heterozygous).

### Segregation distortion and detection of linkage

We developed and implemented in MapDisto a new approach to identify linkage groups (LG) in case of segregation distortion (SD). Several patterns of SD can occur and will lead to a variety of possible effects on the sensitivity and specificity of statistical tests for detection of linkage. The method is detailed in the joint Appendix.

To illustrate the usefulness of the new method, we discuss the treatment of a dataset consisting of 44,398 SNP markers detected by the GBS method described in (Heffelfinger et al. 2014), in the rice population IR64 x Azucena made of 181 RILs and developed at the IRD (Montpellier, France) (Fragoso et al. in preparation). Severe SD occurs on several chromosomes in this population, due to the action of pollen and embryo sac sterility genes. This is commonly observed in *O. sativa* ssp. *indica* x ssp. *japonica* crosses (Harushima et al. 1996). We first tried the classical LOD score method. In order to find LGs corresponding to the twelve rice chromosomes, we had to raise the LOD score threshold to 9.0. However, this caused some LG to be cut into several LGs, because of remaining gaps in the map. We then tried the independence Chi-squared (*χ*^2^_i_) method, designed to improve detection of linkage in case of SD under certain conditions (Lorieux et al. 1995a). This method allowed separating all the LGs with a *χ*^2^_i_ statistic threshold equivalent to LOD = 6.0, except for chromosomes 4 and 7 that remained linked at a *χ*^2^_i_ statistic equivalent to LOD = 9.0. The reason for this is the type of SD occurring at these loci, that is, selection affected only one of the four genotypic classes instead of selection on alleles. With the new method, all the 12 groups were correctly separated at an equivalent to LOD = 5.2 (see Appendix).

### Data imputation and correction

MapDisto employs the new LB-Impute (for Low coverage, Biallelic Impute) algorithm to impute data in biallelic populations with residual heterozygosity typed via low-coverage sequencing (Fragoso et al. 2016). This usually refers to F_2_ or BC_1_F_1_ populations generated from inbred parents, though any population originating from the selfing of a single F_1_ is compatible. LB-Impute is capable of both correcting erroneous genotypes, such as false homozygotes caused by low read coverage, and imputing missing data. The algorithm resolves missing and erroneous genotypes in these populations via a Hidden Markov Model (HMM). In this HMM, emission probabilities are the binomial probabilities of emitting a parental genotype based on the depth of read coverage. Transition probabilities, which represent recombination, are treated as a function of physical distance. Double recombination events may be treated as two independent transition events or a single one depending on the type of population.

Using the emission and transition probabilities, the Viterbi algorithm (Viterbi 1967) is applied from each end of the chromosomes to calculate the most likely sequence of parental states. The consensus sequence of parental states is chosen. Missing markers flanked by the same parental state are assigned that parental state. Moreover, a parental state may be assigned to a genotype that differs from the parental genotype, resulting in a corrected genotype. If the two paths do not agree on a state, however, that marker is left as missing. This often results in missing markers adjacent to recombination breakpoints. Additionally, in order to increase accuracy, LB-Impute has the capability to “look ahead,” or continue to calculate the path past a given marker, to measure how a certain parental state may influence future path decisions. This increases accuracy, but also increases runtime.

LB-Impute is featured as a command in this release of MapDisto to impute genotype datasets. More specifically, LB-Impute is compatible with VCF v4.1 files and produces a new v.4.1 VCF file as output. This new file can be loaded into MapDisto via the VCF conversion tool or analyzed independently. Pseudo-multithreading is obtained by splitting the VCF into separate chromosomes (available for Apple OS X only).

As discussed above, it is likely that variants adjacent to recombination breakpoints will remain as missing after LB-Impute imputation. Although LB-Impute produces a set of high-confidence imputed genotypes, missing markers frustrate trait mapping and genetic map calculation; thus requiring further imputation of these ambiguous intervals. For this reason, a breakpoint imputation algorithm, BP-Impute, was developed in R (manuscript in preparation) and is now available as a command in MapDisto. BP-Impute, also a HMM based method, works by placing two Markov chains on an ambiguous, missing interval. Each chain moves in either direction over the interval, and is constrained to the parental state of the left or right-flanking LB-Impute variant. The transition probability is the proportion of recombinant individuals over the interval, and the emission probabilities are binomial read coverage probabilities just as in LB-Impute. Therefore, each chain represents the probability of remaining in the constrained parental state. The pair of probabilities at each missing variant are normalized to be no greater than 1 and are then taken as a weighted average. These probabilities may be assigned discrete genotypes by applying least squares, using the Assign Genotypes command, which runs an R script that converts genotype probabilities to discrete values. The result is a complete genotyping dataset, with no missing data, suitable for trait mapping or calculating genetic maps. Future improvements to BP-Impute will include improved transition probability estimation (through the use of population-based local recombination rates) and relaxing of the assumption of only one recombination event in an ambiguous interval (which underestimates the genetic map).

### QTL search

Although MapDisto already has built-in features to perform F-test and distribution-free Kruskal-Wallis quantitative trait locus (QTL) search, this new version now provides a graphical interface to perform more advanced analyses, empowered by the R/qtl package (Arends et al. 2010). It provides access to the ‘onescan’ and ‘twoscan’ (for two-dimensional interaction analysis) scan types using different methods: interval mapping, Haley & Knott, extended Haley & Knott, simple marker regression. The user can choose to work on all traits or a specific one, and to all chromosomes or a specific one. Graphical results (one and two-dimension scan, trait summary) are displayed as Portable document format (PDF) files. Also, one of the most tedious tasks in using R/qtl is preparation of input data files; MapDisto performs transparent, automatic formatting and exporting of input data files as R/qtl-compatible files.

## General improvements

The new version has an improved, clearer interface. Use of the Java modules and R/qtl is transparent and doesn’t require any programming or command line ability.

Many functions in the VBA module were re-written and now execute much faster compared to v.1.7. The new version also corrects several bugs found in v.1.7.

## Configuration requirements and installation

### Microsoft Windows platform

MapDisto runs within Excel 2010 or later (Excel 2007 is no longer supported, although still compatible). Windows 7 or later is recommended.

### Apple OS X platform

MapDisto v. 2.0 runs within Microsoft Excel 2011, on any version of OS X from 10.6 (Snow Leopard) to 10.12 (Sierra). An experimental version for the 64-bit Excel 2016 is also available.

To handle very big datasets — typically > 45,000 SNPs —, it is strongly recommended to use the 64-bit versions of both the operating system and Excel. The Apple OS X system is 64-bit by default, as is the last Excel 2016 update. Windows users need to install specific 64-bit versions of both Windows and Excel.

The minimum recommended hardware configuration is a microcomputer with at least 4 GB of RAM and a recent microprocessor. Users should make sure than there is sufficient available RAM for Excel.

The “MapDistoAddons.jar” and “LB-Impute.jar” modules Java modules are seamlessly installed at the first launch of the program. They need a Java Runtime Environment (Windows: version 7 or later; OS X: version 1.7 or later) to run, which either comes pre-installed with the operating system or is downloadable free of charge from the Oracle downloads pages (https://www.oracle.com/java).

For R/qtl commands, the free R statistical package (https://www.r-project.org) is required. The R/qtl library is automatically installed by a command of the MapDisto R/qtl interface.

In order to work properly, MapDisto should be placed in any subfolder in the users’ “Documents” folder. Windows users should install R directly at the root of the C: hard drive.

## Acknowledgements

This study was supported in part by NSF Awards 1444478 and 1419501, and by the Biomedical Informatics Research Training at Yale, project T15 LM 007056 (co-Directors of the Grant Cynthia Brandt and Michael Krauthammer). Computational analyses were performed on the Yale University Biomedical High Performance Computing Cluster, which is supported by NIH grants RR19895 and RR029676-01.

## Appendix: Building linkage groups in case of segregation distortion

We describe the case of a RIL population derived from pure parental lines. We consider only homozygous loci of allele A (parent 1) or B (parent 2). Considering two loci that are to be tested for linkage, there are four possible genotypes in the progeny: AA, AB, BA, and BB, and the observed counts in the contingency table are noted *a*, *b*, *c* and *d*, respectively.

Let’s consider the case of selection against one of the two recombinant genotypes, *e.g*. class *b*. In this case, the commonly used linkage-or independence-Chi-squared tests for linkage are biased and a new linkage test has to be derived. It is easy to show that, in this case, the maximum-likelihood (ML] recombination fraction in the F_1_ is:

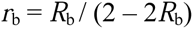

where *R*_b_ = 2*c* / (*a* + 2*c* + *d*)

The corresponding *χ*^2^ statistic is:

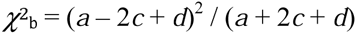

It has one degree of freedom (d.o.f.].

This test is calculated over the population size *n’*= *a* + 2*c*+ *d*. Note that *n’* can be significantly different from *n* = *a* + *b* + *c* + *d*, so we use the correcting factor *n* / *n’* to get a *χ*^2^ value which is relative to *n* (*χ*^2^ is increasing linearly with *n*). This allows normalizing the population size across all tests. So we get:

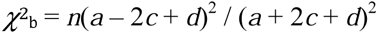

One can derive similar formulas for cases of selection against class *a*, *c* and *d*, so we have the four *χ*^2^ statistics:

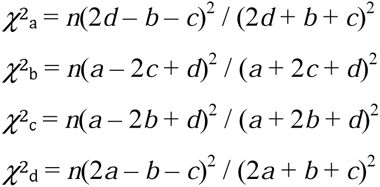

It is desirable to translate these values as LOD score equivalents. This would allow users to directly compare the results of classical LOD score and *χ*^2^ tests, and to compare the same threshold values for tests that take SD into account and tests that do not.

To derive the LOD score equivalents, let’s consider the following relation:

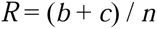

*R* is the observed fraction of recombinant individuals, and the estimated recombination fraction (RF) in the F_1_ is:

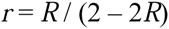

This gives the inverse function:

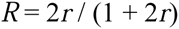

The LOD score for linkage in RILs is:

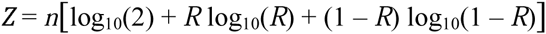

And the Chi-squared for linkage is:

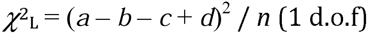

Remark that *a* + *d* = *n*(1 - *R*)and *b* + *c* = *nR*. So *χ*^2^_L_ can be re-written as follows:

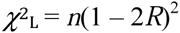

Thus, a LOD score for linkage can be calculated from Chi-squared values using the following formulas:

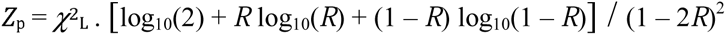

if 0 <= *R* < 0.5.

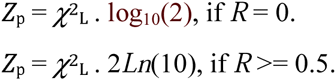

We can now develop the algorithm for detecting linkage in case of segregation distortion in RILs.

## Algorithm

First, compute the observed fraction of recombinant individuals *R*= (*b* + *c*) / *n*. Then, consider the three possible cases:

***Case 1*: *R* = 0**

- The F_1_ recombination fraction is *r* = 0.

- The LOD score is *Z* = *n*log_10_(2).

- End of the algorithm.

***Case 2*: *R* > 0.5**

- Set *R* to 0.5.

- The F_1_ recombination fraction is *r* = 0.5.

- The LOD score *Z* is set to 0.

- End of the algorithm.

***Case3*: 0 < *R* <= 0.5**

- Check segregation of locus 1 and locus 2 genotypic frequencies for Mendelian expectations. The test for Mendelian segregation of a locus is:

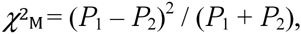

where *P*_1_ and *P*_2_ are the counts of allele 1 and allele 2, respectively.

The probability *p* of rejecting Ho: *P*_1_ = *P*_2_ is calculated with 1 d.o.f.

*Note*: remark that *P*_1_ and *P*_2_ cannot be extracted from *a*, *b*, *c*, *d* classes when there is missing data.

***Case 3.1*: None of the two loci shows Mendelian segregation**

- The F_1_ recombination fraction is:

Backcross or doubled haploid: *r* = *R*

RIL: *r* =*R* / (2 – 2*R*)

-The LOD score is:

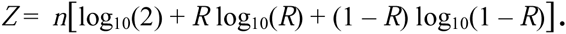

- End of the algorithm.

***Case 3.2*: At least one locus shows segregation distortion^2^**

1 - Compute the six Chi-squared tests (*χ*^2^_L_, *χ*^2^_i_, *χ*^2^_a_, *χ*^2^_b_, *χ*^2^_c_, *χ*^2^_d_). The independence Chi-squared testis:

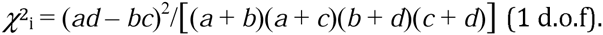

- Retain *χ*^2^_a_ if *a* < *d* AND *χ*^2^_a:d_ is significant (e.g. at 5%),
where *χ*^2^_a:d_ = (*a* – *d*)^2^ / (*a* + *d*) (1 d.o.f).

- Retain *χ*^2^_b_ if *b* < *c* AND *b* < *a* AND *b* < *d* (*b* is the smallest class) AND *χ*^2^_b:c_ is significant (e.g. at 5%),
where *χ*^2^_b:c_ = (*b* – *c*)^2^ / (*b* + *c*) (1 d.o.f).

- Retain *χ*^2^_c_ if *c* < *b* AND *c* < *a* AND *c* < *d* (*c* is the smallest class) AND *χ*^2^_b:c_ is significant (e.g. at 5%).

- Retain *χ*^2^_d_ if *d* < *a* AND *χ*^2^_a:d_ is significant (e.g. at 5%).

2 - Determine the least of the retained tests.

3 - Only for the least *χ*^2^, re-compute *R* using the following rules^3^:

- If *χ*^2^_L_ is least, then *R* = (*b* + *c*) /*n*.

- If *χ*^2^_i_ is least, then *R* = 0.5 - (*χ*^2^_i_ / 4*n*)^0.5^.

- If *χ*^2^_a_ is least, then *R* = (*b* + *c*) / (*b* + *c* + 2*d*).

- If *χ*^2^_b_ is least, then *R* =2*c* / (*a* + 2*c* + *d*).

- If *χ*^2^_c_ is least, then *R* = 2*b* / (*a* + 2*b* + *d*).

- If *χ*^2^_d_ is least, then *R* = (*b* + *c*) / (2*a* + *b* + *c*).

4 - Compute the LOD score equivalent:

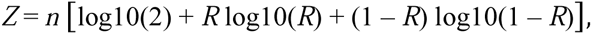

where *R* is replaced by its computed value in step 3.

5 - Compute the F_1_ recombination fraction:

Backcross or doubled haploid: *r* = *R*

RIL-SSD: *r* =*R* / (2 – 2*R*)

- End of the algorithm.

We can then use *r* and *Z* values to test for linkage in case of segregation distortion.

## Footnotes

1 See http://mapdisto.free.fr/MapDisto/Refs for a list of references that use MapDisto.

2 We know that, in the case where only one locus’ segregation is distorted, the classical estimate of r is unbiased (Lorieux *et al.* 1995a). Thus, Case 3.2 theoretically reduces to the case where the two loci show SD. However, situations where locus 1 is severely distorted and locus 2 shows SD just under the significance level can occur. In such situations, the classical estimate for *r* is biased. We thus take a conservative approach where the hereby algorithm is applied to any situation where at least one of the two loci shows SD.

3 The idea behind choosing the least of the six Chi-squared tests is that the smallest Chi-squared has the best fit with the data.

